# Exploring neural manifolds across a wide range of intrinsic dimensions

**DOI:** 10.1101/2025.07.01.662533

**Authors:** Jacopo Fadanni, Rosalba Pacelli, Alberto Zucchetta, Pietro Rotondo, Michele Allegra

**Author notes:** Electronic address.

## Abstract

Recent technical breakthroughs have enabled a rapid surge in the number of neurons that can be simultaneously recorded, calling for the development of robust methods to investigate neural activity at a population level. In this context, it is becoming increasingly important to characterize the neural activity manifold, the set of configurations visited by the network within the space defined by the instantaneous firing rates of all neurons. The intrinsic dimension (ID) of the manifold is a key parameter allowing to relate neural trajectories with the ongoing network computations. While several studies suggested that the ID may be typically low in neural manifolds, contrasting findings have disputed this statement, leading to a wide debate. Part of the disagreement may stem from the lack of a shared and robust methodology to measure the ID. In the case of curvature, linear methods tend to overestimate the ID; in the case of undersampling, nonlinear methods tend underestimate it. Here we show that adapting the full correlation integral (FCI) method yields an estimator that is robust to both curvature and undersampling. We tested our metric on artificial data, including neural trajectories generated recurrent neural networks (RNNs) performing simple tasks and a benchmark dataset consisting of non-linearly embedded high-dimensional data. Our methodology provides a reliable and versatile tool for the analysis of neural geometry.

## I. INTRODUCTION

Traditionally, brain circuits were analyzed by characterizing the functional role of individual units. The advent of large-scale neuronal recordings [1] opened the door to a different approach, focused on understanding the collective activity of all units in terms of representations and computation within abstract spaces [2, 3]. Central to this perspective is the concept of *neural manifold* [4, 5], the set of configurations visited by the system within the space defined by the activity of all units [6–9]. Visualizing, analyzing, and modeling neural manifolds requires identifying a minimum set of coordinates describing the manifold. The minimum number of coordinates is called the manifold’s *intrinsic dimension (ID)*. The ID encapsulates the complexity of neuronal dynamics, in the classical sense of the relation between the individual units and the whole system [10]. The ID also bounds the possible number of cognitively and behaviorally relevant variables directly encoded by neurons at a population level.

Characterizing the ID of neural manifolds is essential to understand how the collective neuronal activity supports the execution of cognitive tasks. Several studies investigating neural manifolds during simple instructed tasks reported a low (≲ 10) ID [11–16], inspiring a large body of theoretical work aiming to explain how low-dimensional activity can emerge from structural constraints [17–20]. However, many other studies reported a high ID, typically in presence of complex stimulus spaces and/or loose task structure [21–24]. A thorough comprehension of what determines the geometry of neural manifolds, of which the ID is a key characteristic, is still missing. The specific type of computation or ‘task’ being performed is certainly a crucial factor, but not the unique one: for any given task, neural networks may also need to strike an optimal trade-off between conflicting demands such as robustness (which favors low dimensional codes [23]) and ease of readout (which favours high-dimensional codes[25–30]). A clearer understanding demands progress not only in neuronal recordings, but also in analysis methodology. In this context, a key gap is the lack of a reliable and shared methodology to measure the ID of neural manifolds [31].

Research on neural manifolds often emplys linear ID estimation methods [13, 14, 19, 21, 22, 24, 26, 27, 32–39], which are based on the spectrum of the covariance matrix and use several criteria to select the number of ‘relevant’ or ‘significant’ dimensions. ID estimates by linear methods generally coincide with the dimension of the (minimal) linear subspace encompassing the manifold [40], rather than the manifold’s actual ID. As a resut, they can significantly over-estimate the ID in the case of non-linear, curved manifolds, despite the fact that the latter are quite common [41, 42]. An alternative to linear methods is provided by geometric ID estimators [43–45], which are ultimately based on the principle that distances in the data follow scaling laws depending parametrically on the ID. The major shortcoming of these methods is that they are not robust to undersampling, requiring a sample size growing exponentially with the ID. As a consequence, they are generally unable to yield proper ID estimates when the ID is large (≳ 10). These shortcomings were highlighted in Ref. [31], where simulated data were used to show that no common method can correctly identify the ID of non-linear, high-dimensional neural manifolds.

Here, we propose a pipeline to evaluate the ID of neural manifolds by leveraging the Full Correlation Integral (FCI) estimator [46]. The major advantage of FCI is its remarkable robustness with respect to undersampling, as it can estimate high IDs from samples whose size scales only linearly with the ID. On the other hand, FCI typically over-estimates the ID in presence of curvature, similarly to linear methods. To overcome this drawback, our pipeline employs a local version of FCI, called *local FCI (lFCI)*. We test lFCI on two sets of artificial data: (i) neural manifolds generated by recurrent neural networks (RNN) trained on the ‘cog-task’ battery [47]. This task set, inspired by real tasks used in neurophysiological studies on non-human animals, involves basic cognitive processes such as working memory, inhibition, and context-dependent integration. For this reason, it has later been considered in many studies, becoming a sort of golden standard [48–52]. The simplicity of the stimulus-response patterns in RNNs reflects into a strong and consistent expectation about the low dimensionality of the corresponding neural manifolds, which is not generally grasped by commonly-used ID estimators. (ii) a benchmark dataset of synthetic neural trajectories, embedded in high-dimensional space both linearly and non-linearly [31]. Trajectories are generated from an empirical distribution of firing rates, obtained from multi-electrode array recordings in the macaque primary motor cortex. As shown by Altan et al., common linear and nonlinear estimators fail in the high-dimensional case, underestimating the dimension as soon as *ID ≳* 10. For the cog-task battery, we find that our estimator works much better than linear estimators and several geometric estimators, providing consistent estimates for different trainings of the networks. For the benchmark dataset, our estimator performs comparably with the best linear estimator (in the linear embedding case) and performs better than other methods (in the non-linear case).

This manuscript is organized as follows: in Methods, we provide a review of state-of-the-art ID estimators, including FCI, and illustrate the process to generate the synthetic datasets investigated. In the Results section, we present our pipeline (lFCI), and the results of its application to synthetic neural manifolds.

## II. METHODS

### A. Intrinsic Dimension Estimation

Consider a set *X* = {**x**_1_, …, **x**_*P*_} of *P* points embedded in an ambient space of dimension *N*. Assuming that the points are contained, exactly or approximately, within a manifold or hypersurface of *intrinsic dimension D < N*, one can try to estimate *D*. There exists a wide array of ID estimation methods (for a classic review, see [53]). To be suitable for characterizing neural activity manifolds, an estimator should ideally work well for non-linear, curved manifolds and a large interval of dimensions, including high dimensions where the manifold is undersampled.

Linear ID estimation methods are based on the spectrum of the covariance matrix of the points. Let *λ*_*k*_, *k* = 1, …, *N* be the eigenvalues of the covariance matrix in decreasing order (*λ*_1_ ≥*λ*_2_ ≥ …≥ *λ*_*N*_). The corresponding eigenvectors are commonly termed *principal components (PCs)* [54]. The ID can be determined by assessing the ‘relevant number’ of PCs. The most common method consists in searching for a linear subspace accounting for a large fraction of the total variance 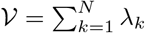, i.e., identifying the minimum *K* such that

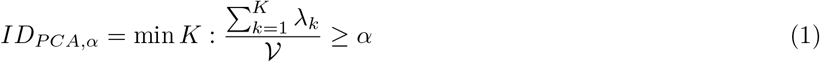

(typical choices are *α* = 0.8, 0.9, 0.95, 0.99). Another method computes the *participation ratio* [55]

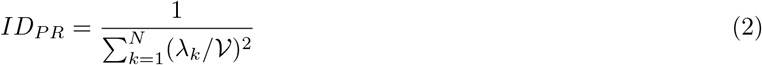

*ID*_*PR*_ corresponds to an ‘effective number’ of relevant dimensions, as typically *ID*_*PR*_ ≃ *e*^*H*(***λ***)^ with 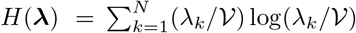 the entropy of the covariance matrix spectrum. Finally, *parallel analysis (PA)* [56] determines the number of significant PCs through a null model. Surrogate data are constructed by resampling the original data by shuffling each data coordinate independently. For each surrogate dataset, the eigenvalues of the covariance matrix are computed. By considering a large number of shuffling, one obtains a null distribution of eigenvalues. Let *ν*_*α*_ be the (1 −*α*) · 100-th percentile of the null distribution (typically, *α* = 0.05). One finally considers as significant all original eigenvalues exceeding this critical value,

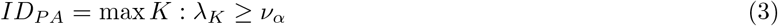

Linear methods are appropriate only when the manifold is flat, i.e., it lies on a hyperplane. In presence of curvature, they can largely overestimate the ID.

Among non-linear methods, the most commonly used are *fractal* [57, 58] and *nearest-neighbor* methods [43–45]. Fractal methods are based on the fact that the number of points found within a spherical volume of radius *r* follows an ID-dependent scaling law. The most common example is the Correlation Dimension (CorrDim) method first proposed in the classic paper by Grassberger and Procaccia [57]. CorrDim focuses on the *correlation integral* (the density of points within a given cutoff distance *r*):

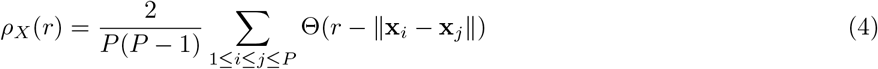

If the points are sampled from a distribution on a *D*-dimensional manifold, then *ρ*_*X*_ (*r*) ~*r*^*D*^ for *r* → 0. Thus, one can extract the ID by measuring the slope of the linear fit of *ρ* as a function of *r* in a log-log scale (*D* = lim_*r*→0_ log *ρ*_*X*_ (*r*)*/* log *r*). This method can work reasonably well for non-linear manifolds, but is fragile with respect to undersampling, as it requires a number of points scaling exponentially with the ID (“curse of dimensionality”), and thus underestimates high IDs. Nearest-neighbor methods are based on the fact that distances between nearest neighbors obey statistical relations that depend parametrically on the ID. The max-likelihood estimator [43] assumes that the data are distributed on a *D*-dimensional manifold and that the density is uniform in a neighborhood containing the first *K* neighbors of each point. Under this assumption, consider 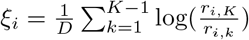 where *r*_*i,k*_ are the distances of the *k*-th neighbors of point **x**_*i*_ in the data. One can show that

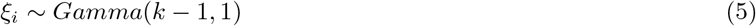

from which one can estimate *D* with a maximum likelihood approach on the empirical *ξ*_*i*_. The two-nearest-neighbor (Two-NN) estimator [45] restricts the uniform density assumption to the neighborhood containing only the first 2 neighbors of each point. Under this assumption, one can show that

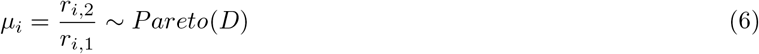

and one can infer *D* through a non-linear fit of the empirical distribution of *µ*_*i*_. Also nearest neighbor methods incur the curse of dimensionality.

### B. The FCI estimator

The full correlation integral (FCI) estimator proposed by Erba et al. [46] was shown to be remarkably robust to undersampling, being able to correctly estimate large IDs even in the extreme case *P < D*. The starting point of the method is provided by CorrDim. To address the curse of dimensionality, Erba et al. made the more restrictive assumption that points are sampled from a rotationally-invariant probability distribution on a *D*-dimensional hyperplane. In this case, after normalization of each point to unit norm, the data must lie on a *D* − 1-dimensional hypersphere (e.g., a circle for *D* = 2, a spherical surface for *D* = 3), and one can exactly calculate the average correlation integral as a function of *r*:

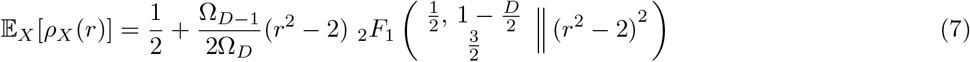

where _2_*F*_1_ is the (2, 1)-hypergeometric function and Ω_*D*_ is the solid angle in dimension *D*. To stress that this analytic formula holds away from the small radius limit of CorrDim, the average correlation integral is named *Full Correlation Integral (FCI)*. With this definition, the FCI, as a function of *r*, has a sigmoidal shape with a slope that depends on *D*. Performing a non-linear regression on the empirical density of neighbors (4) using the FCI equation (7) results in an algorithm to determine the ID, shown in Fig. 1. In detail, the steps are:

**FIG. 1:**
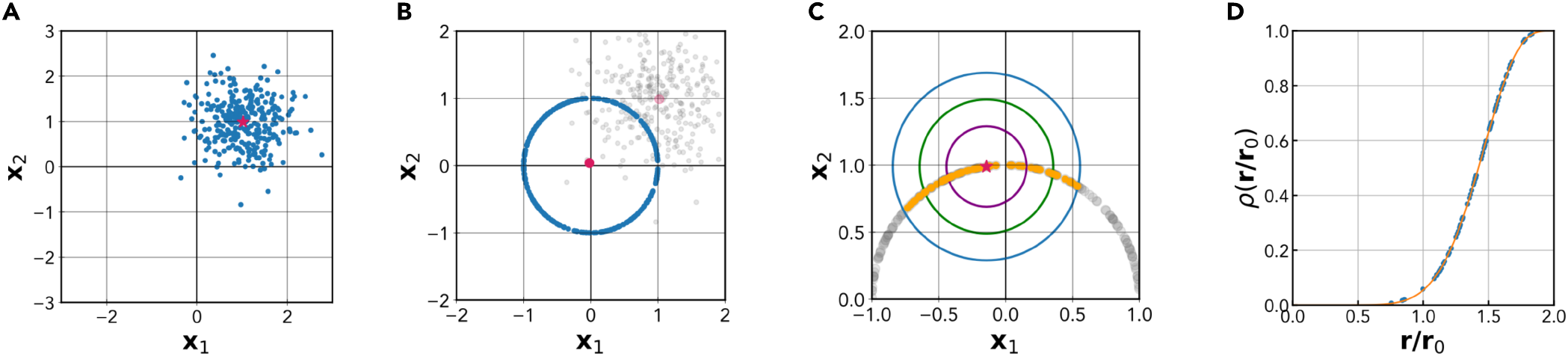
The Full Correlation Integral (FCI) method. Panels A-D illustrate the basic steps of the FCI method for ID estimation. **A** The original *D*-dimensional dataset (the red dot indicates the barycenter). **B** Centering and normalization of the dataset: the data points are projected on an unitary sphere in *D* −1 dimensions. **C** The empirical correlation integral of the dataset, *ρ*(*r/r*_0_), is measured as a function of the normalized radius *r/r*_0_. This amounts to computing how many pairs of points are within a distance *r*. **D** ID estimation: *ρ*(*r*) is fitted with its theoretical model given by Eq. (7), which depends parametrically on the ID. To obtain the true ID, the estimated ID is increased by one, compensating the removal of one degree of freedom during the normalization step.

1. *Center and normalize the data* (Fig. 1A,1B). Calculate the center of mass **b** of the empirical data as 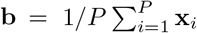 and subtract this quantity from each data point, obtaining a data set centered at the origin. Normalize each sample as **x**_*i*_ = **x**_*i*_*/*||**x**_*i*_||.
2. *Fit the FCI*. Measure the empirical correlation integral of the dataset at different radii *r* (Fig. 1C) and perform a non-linear regression of this empirical neighbors density using the average FCI in Eq. (7) as the nonlinear model, and *D* as the free parameter (Fig. 1D); increase the estimate by one, *D* → *D* + 1 to reinstate the degree of freedom removed during the normalization step.

Strong deviations from the predicted functional form of the FCI can be detected by the goodness of fit (GoF), measured as the root-mean-squared deviation of the empirical FCI from its best fit using Eq. (7). Empirically, a bad fit is obtained when GoF ≳ 0.02 (in Fig. S1, we report examples of fits with different values of GoF). By construction, the FCI method is exact for linearly embedded Euclidean spaces sampled with a rotationally-invariant probability distribution. Erba et al. showed that the method is remarkably robust with respect to violations of the rotational invariance assumption. In particular, they showed that FCI estimates work very well even in some cases of strongly non-isotropic distributions (e.g., points sampled from corners of *D*-dimensional hypercubes). In addition, estimates are very robust with respect to undersampling, as correct estimates of *D* ~ 200 can be obtained from as little as *P* ~ 20 points.

### C. Recurrent Neural Networks performing Cog-Tasks

RNNs have become a customary tool of computational neuroscience research [59–62]. We trained RNNs to solve the ‘cog-task’ battery first introduced by Yang et al. [47]. For simplicity, we maintained the same RNN architecture and dynamics as originally proposed.

The network has *N*_*rec*_ = 256 recurrent units in the hidden layer coupled with *N*_*in*_ = 65 input units and *N*_*out*_ = 33 output units. The input layer is composed of one fixation unit and two rings of 32 units representing two stimuli. The output layer has one fixation unit and a ring of 32 units representing motor output.

In the rate model of network activity, each neuron is represented by a continuous variable 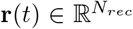 representing the neuron’s membrane potential. The dynamics are described by the equation:

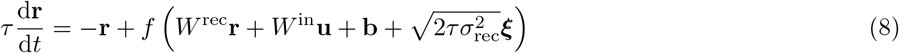

Here, *τ* = 100 ms represents the neuronal time constant, reflecting the slow synaptic dynamics driven by the NMDA receptors. The variable 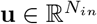 denotes input to the network, while 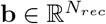 is the bias or background input. The function *f* (·) captures the neuronal nonlinearity, and ***ξ*** represents *N*_rec_ independent Gaussian white noise processes with zero mean and unit variance, and *σ*_rec_ = 0.05 quantifies the noise intensity. A typical Softplus function is used, *f* (*x*) = log (1 + exp (*x*)), which after re-parametrization is very similar to the neuronal nonlinearity (frequency-current curve) commonly used in neural circuit models. A set of output units **z** read out nonlinearly from the network,

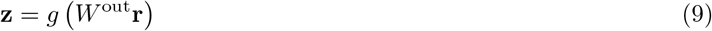

where *g*(*x*) = 1*/*(1 + *exp*(−*x*)) is the logistic function that limits output activities between 0 and 1. *W*^*in*^, *W*^*rec*^, *W*^*out*^ are the input, recurrent, and output connection matrices, respectively. After using the first-order Euler approximation with a time-discretization step Δ*t*, we have

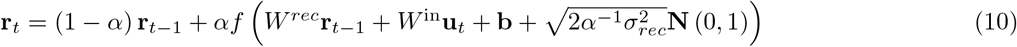

Here, *α* ≡Δ*t/τ*, and *N* (0, 1) represent the standard normal distribution. We use a discretization step Δ*t* = 20 ms. We imposed no constraint on the sign or structure of the weight matrices *W*^*in*^, *W*^*rec*^, *W*^*out*^. The network and training were implemented in TensorFlow[63]. The network receives three types of noisy input,

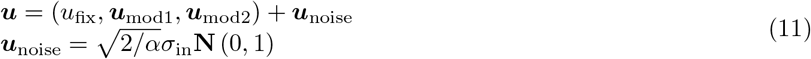

The input noise strength is set to *σ*_*in*_ = 0.01. The fixation input *u*_fix_ is usually at the high value of 1 when the network is expected to maintain fixation, and changes to 0 when the network is supposed to react. The stimulus inputs **u**_mod1_ and **u**_mod2_ represent one-dimensional circular variables characterized by the angle around a circle. This mirrors what happens in typical experiments with primates, involving *directional stimuli and responses* (for instance, the stimulus is a cloud of dots moving in a given direction, and the animal is asked to move the arm in the same direction). There are 32 units in each of the corresponding stimulus rings, with their preferred directions evenly distributed from 0 to 2*π* (*ψ*_*i*_ = 2*π* · *i/*32 = *π/*16 · *i*). For unit *i*, which has a preferred direction *ψ*_*i*_, its activity in response to a stimulus at direction *ψ* is given by

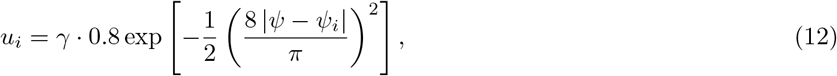

where *γ* represents the stimulus intensity. For multiple stimuli, input activities are summed.

The tasks used here are the same as those used by Yang et al.[47]. They selected 20 interrelated tasks, representing common tasks used in neurophysiological studies on nonhuman animals, and useful to understanding basic cognitive processes such as memory-guided response, simple perceptual decision making (DM), context-dependent DM, multisensory integration, parametric working memory, inhibitory control, delayed match-to-sample and delayed match-to-category tasks. The tasks can be divided into three families: the Go, Decision Making (DM), and Matching families.

a. *Go family*. In this family, a single stimulus is randomly shown either in modality 1 or 2, and the response should be in the same direction of the stimulus. This family includes the forward-Go (Fd-Go), in which the network should respond when the fixation cue goes off; the reaction time Go (RT-Go), in which the network should respond as soon as the stimulus appears; the delay go (Dly-Go), in which the stimulus is turned off before the go and the network has to respond only after the fixation cue goes off. In the ‘anti’ versions of the tasks (Fd-Anti, RT-Anti and Dly-Anti), the network should respond in a direction opposite to that of the stimulus.
b. *DM family*. In this family, two stimuli are shown simultaneously and are presented till the end of the task. A stimulus is drawn randomly on the whole circle while the other is drawn uniformly between 90° and 270° away from the other. The network has to respond in the strongest of the two directions. The family includes DM1 and DM2 where the two stimuli are presented only in modality 1 (respectively, 2); Ctx DM1 and Ctx DM2 in which the network has to consider only the two stimuli presented in modality 1(2) ignoring those in the other modality; MultiSensory DM in which the network has to respond in the direction of the stimulus that has a stronger combined strength in modalities 1 and 2. In the delayed versions of the tasks (Dly DM 1, Dly DM 2, Ctx Dly DM 1 and Ctx Dly DM 2), the two stimuli are not contiguous in time. They are both shown briefly and are separated by a delay period. Another short delay period follows the offset of the second stimulus before the response.
c. *Matching family*. In this family, two stimuli are presented consecutively in modality 1 or 2 and separated by a delay period. The response depends on whether or not the two stimuli “match”. This family includes the delayed-match-to-sample (DMS), delayed-non-match-to-sample DNMS, delayed-match-to-category (DMC) and delayed-non-match-to-category DNMC tasks. In DMS and DNMS, two stimuli match if they point towards the same direction; in DMC and DNMC, if their direction belongs to the same ‘category’, the first category ranging from 0 to 180°, the other from 180 to 360°. In DMS and DMC the network has to respond towards the second stimulus when there is a match, and fixate otherwise; in DNMC and DNMS, it has to respond towards the second stimulus when the two stimuli are not matched, and fixate otherwise.

### D. Synthetic neural recordings

Altan et al.[31] created synthetic data to investigate the performance of ID estimators on neural manifolds. In particular, they constructed synthetic data of known dimensionality, mimicking the properties of multi-electrode neural activity recorded in the primary motor cortex of macaques. Data generation starts by extracting *d* × *M* samples from an empirical distribution of firing rates obtained from multi-electrode array recordings of the macaque primary motor cortex [64] with a binning of 50 *ms*. Samples are extracted randomly across all recorded neurons and time bins, resulting in temporally uncorrelated samples. The *d*-dimensional data are then temporally smoothed (with a Gaussian kernel of s.d. 50 *ms*) and multiplied by a *N* × *d* mixing matrix with entries randomly selected from a Gaussian distribution of zero mean and unit variance. This produces a dataset *X* composed of *M* samples of dimension *N*. This embedding step does not change the intrinsic dimension *d*. The value of *d* ranges from 3 to 40, while *N* = 96 is similar to the number of electrodes in multi-electrode measurements.

In order to produce a dataset with strongly non-linear properties, an exponential nonlinearity is computed on each of the *N* variables in *X*, yielding a data set

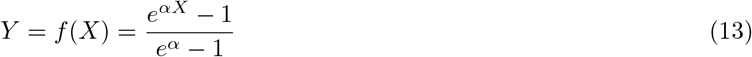

where *α* determines the degree of nonlinearity. Also this non-linear embedding procedure should in principle preserve the ID.

## III. RESULTS

### A. The local FCI pipeline

We start by presenting our pipeline for ID estimation, called the local Full Correlation Integral (lFCI) method. The underlying rationale is exploiting a local version of the FCI estimator (see Methods and [46]) to a achieve a robust ID estimator working across a wide range of dimensions and for curved manifolds. As thoroughly expounded in [46], FCI is robust in high dimension, but it can overestimate the ID in presence of curvature. This can be illustrated with a simple example. On points sampled from a uniform distribution on a two-dimensional plane, FCI yields a very accurate estimate, *D* = 1.97 (Fig. 2A). On points uniformaly sampled on a Swiss roll - a curved two-dimensional surface embedded in three-dimensional space - FCI overestimates the dimension, yielding *D* = 2.73 (Fig. 2E).

**FIG. 2:**
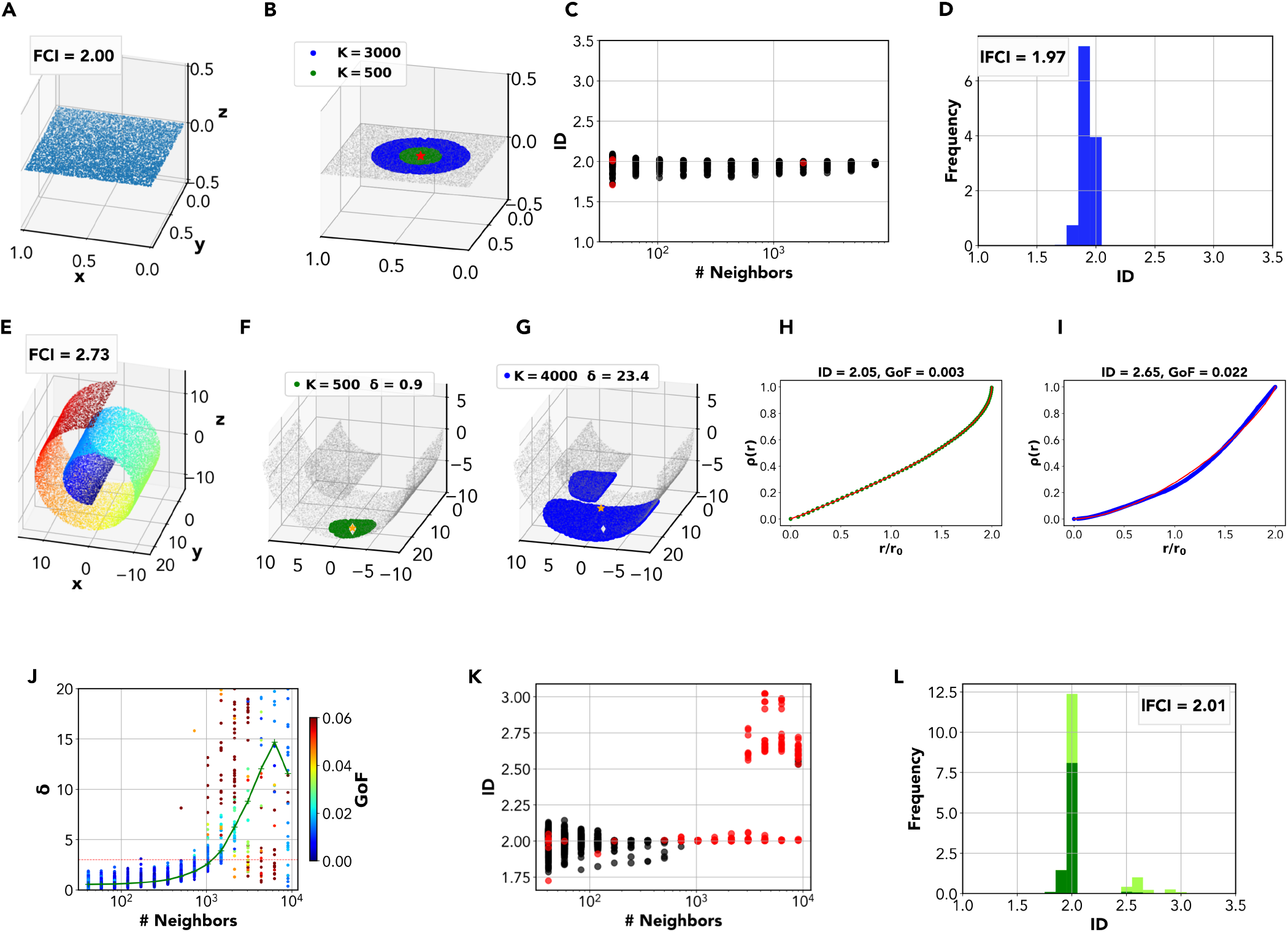
The Local FCI method. **A** A dataset with uniform distribution on a plane, *ID* = 2. FCI correctly identifies the ID of the dataset, yielding *ID*_*FCI*_ = 2.00. **B** Local ID estimates can be computed using FCI on neighborhoods of increasing size *K*. **C** In the *multiscale ID plot*, local ID estimates are shown as function of the neighborhood size *K*. Estimates are colored according to their reliability, as assessed by Goodness-of-Fit (GoF). **D** All reliable local ID estimates for different *K* are collected in the *local ID histogram*. The mode of local ID estimates (the peak of the histogram) yields the a single overall ID estimate. We call this ID estimation method the *local FCI estimation* (lFCI) method. For the 2-D plane, we obtain *ID*_*lFCI*_ = 1.97. **E** A dataset with points uniformly distributed on a Swiss Roll. Even though the ID is 2, FCI gives *ID*_*FCI*_ = 2.97. The global FCI overestimates the ID due to the curvature of the manifold. **F** Neighborhoods of small size (*K ≲* 10^3^ approximate well the local tangent plane to the manifold. The barycenter is located within the neighborhood (low *δ*). **G** As the neighborhood size increases, neighborhoods depart from the local tangent plane, becoming curved and even possibly disconnected. As a result, the barycenter is located outside of the neighborhood (high *δ*). **I** Some neighborhoods (including most curved ones) yield a bad goodness-of-fit of FCI **J** The ratio between distance of the barycenter and the average distance between point greatly increases when the neighborhoods became large highlighting a curvature in the dataset. **K** In the *multiscale ID plot* for the Swiss Roll, we observe a sharp transition: for *K ≲* 10^3^, local ID estimates are close to 2, while for larger *K* they approach 3. However, we can discriminated bona fide estimates (*δ <* 3 and *GoF <* 0.02), shown in black, from unreliable onbes (*δ* > 3 or *GoF* > 0.02), shown in red. For *K ≳* 10^3^, most estimates are unreliable. **L** The local ID histogram for the Swiss Roll shows a prominent peak at ~2. We obtain *ID*_*lFCI*_ = 2.01.

This limitation can be addressed by exploiting *local ID estimates*. For any given point **x**_*i*_ in the dataset, one can consider its first *K* neighbors,

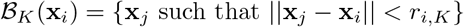

where *r*_*i,K*_ is the distance of the *K*-th neighbor. Local neighborhoods are shown for the plane in Fig. 2B. By restricting the ID estimation to points within the neighborhood ℬ_*K*_(**x**_*i*_), one obtains a local ID estimate *D*_*i,K*_. One can randomly subselect *M* ≫ 1 points and consider neighborhoods of increasing size *K* centered at each of these points, computing *D*_*i,K*_ for each point and neighborhood size. Estimates *D*_*i,K*_ as a function of the scale *K*can be collectively organized in a *multiscale ID plot*. This is shown for the 2-D plane in Fig. 2C: estimates at all scales give values in the narrow range [1.7, 2.2]. By collecting all local estimates for different *K* in a single *local ID histogram*, we obtain a sharp peak around the expected value, 2 (Fig. 2D).

When replicating this procedure on the Swiss roll, we observe that estimates *D*_*i,K*_ remain around 2 for small neighborhood sizes (*K ≲* 10^3^), while they progressively approach the value 3 for larger neighborhoods (Fig. 2K). This is rather unsurprising: for sufficiently small *K*, neighborhoods will be flat, approximating the local tangent plane of the manifold, and local ID estimates will correspond to the manifold’s proper ID; for larger *K*, neighborhoods will become non-flat and estimates will converge to the standard global estimate. This suggests that *a proper ID estimate could be obtained from the collection of local estimates, removing curved neighborhoods from analysis*.

In fact, there are two metrics that can track the departure of local neighborhoods from the tangent plane. The first is simply the FCI Goodness-of-Fit (See Methods). Curved neighborhoods often violate FCI’s core assumption of local isotropy, leading to bad GoFs. The other is the position of the neighborhood’s *barycenter*, 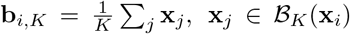. For a flat neighborhood, the baycenter **b**_*i,K*_ lies amidst points of the neighborhood: as a result, the barycenter’s distance from the closest point in the neighborhood will be of the same order of the typical distance between nearest neighbors within the neighborhood. For a curved neighborhood, instead, **b**_*i,K*_ will be displaced from the neighborhood (for an extreme case, think of a spherical surface: its baycenter is the sphere’s center, located far off all surface points). This behavior is effectively captured by the metric

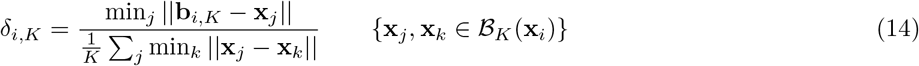

which is simply the distance between the barycenter and the closest neighborhood point, normalized by the average nearest-neighbor distance in **b**_*i,K*_. For flat neighborhoods, we expect *δ* ~ 1, while curved neighborhoods will give *δ* > 1.

The effectiveness of these two metrics can be exemplified by considering two neighborhoods of different sizes on the Swiss Roll. In Fig. 2F, we show a neighborhood with *K* = 500. The neighborhood approximates the tangent plane at **x**_*i*_ and it is nearly flat, as indicated by the value *δ*_*i,K*_ = 0.9. Correspondingly, we obtain a good fit (GoF = 0.003) and a good ID estimate *D*_*i,K*_ = 2.04 (Fig. 2H). In Fig. 2G, we show a larger neighborhood with *K* = 4000. The neighborhood strongly deviates from the tangent plane at **x**_*i*_, as indicated by the value *δ*_*i,K*_ ~ 15. Correspondingly, we obtain a bad fit (GoF = 0.022) and an excessive ID estimate *D*_*i,K*_ = 2.65 (Fig. 2I). The just exemplified behavior is systematic, as shown in Fig. 2J. For small neighborhoods (*K* ≲ 500) we invariably obtain *δ* ~ 1 and GoF *<* 0.02. For larger ones (*K* ≳ 500), *δ* progressively grows and the fit quality degrades.

We consider local ID estimates as *unreliable* whenever fit quality was poor (GoF *<* 0.02) or the local neighborhood could be identified as curved with good confidence (*δ* > 3). Accordingly, in the multiscale ID plot (Fig. 1K) we display reliable estimates in black and unreliable ones in red. Nearly all reliable ID estimates are around 2, while excessive estimates are nearly always spotted as unreliable. While the local ID histogram is bimodal, with a peak at 2 (determined by small neighborhoods) and a peak at ~3 (determined by large neighborhoods), excluding unreliable estimates suppresses the peak at 3 (Fig. 2L).

We thus propose the following *local FCI (lFCI)* pipeline:

1. compute local estimates *D*_*i,K*_ on randomly selected neighborhoods with different values of *K*.
2. Discard estimates corresponding to low fit quality (GoF > 0.02) or large curvature (*δ*_*i,K*_ > 3)
3. Collect all local ID estimates in a histogram and take its highest peak *D*^*\**^ as the best estimate of the overall ID.

This pipeline is expected to work well for sufficiently well-sampled Riemannian manifolds, such that local neighborhoods will approximate an isotropic sampling on the manifold’s tangent plane.

### B. ID of neural manifolds of recurrent neural networks performing the Cog-Task battery

We trained recurrent neural networks (RNNs) to perform the ‘cog-tasks’ battery [47] (Fig. 3A). For each task, we trained *M* = 10 networks. After training, each RNN performed *N* = 200 trials of the task. For each network, we defined the neural manifold as the set of configurations (in the space of activities of the recurrent layer neurons) visited by the network across the *N* trials. We characterized each manifold with common ID estimators.

**FIG. 3:**
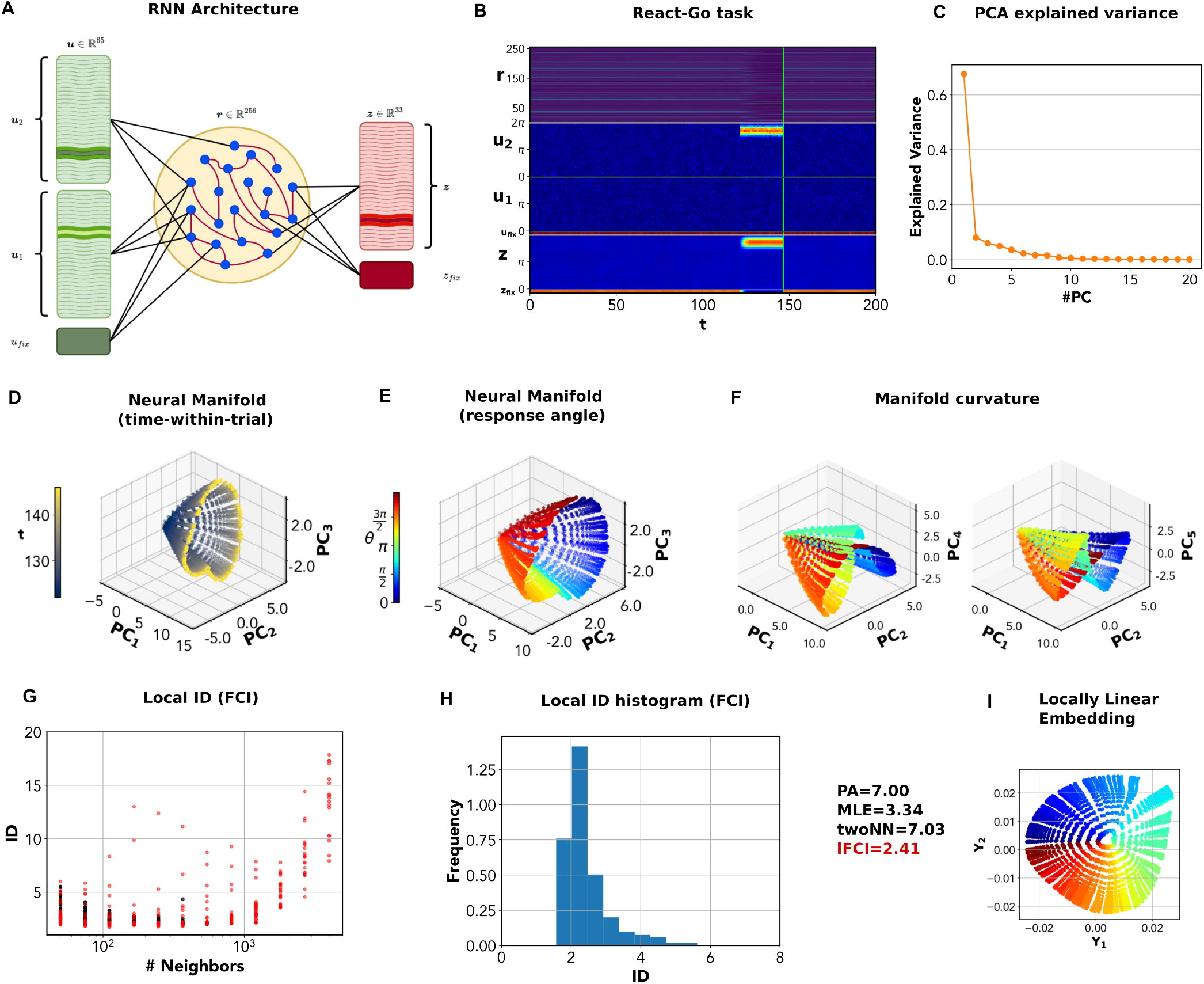
Intrinsic dimension of the neural manifold in the RT-Go task. **A** Schematic structure of the RNN used for the Cog-Task battery. The network has 65 input units: two sets of 32 channels each, *u*_1_, *u*_2_, respectively encode two angular stimuli that jointly determine the network’s response; the fixation input *u*_*fix*_ instructs the network on when to respond. The recurrent layer has 256 units. The output layer has 33 units: 32 units represent an (angular) motor response, *z*, while the last unit represents a ‘fixation’ response *z*_*fix*_. **B** Input, output and network activity for the RT-Go task. In this task, a single angular stimulus is presented, and the network should immediately give a response in the same direction. **C** Variance explained by successive principal components (PCs) of the network activity. **D** Projection of the network activity on the first three PCs. The color code stands for time-within-trial (time from trial onset). *PC*_1_ is nearly aligned with time-within-trial. Starting from the origin (null activity), the trajectories reach a ‘ring’ of attractor points (yellow). **E** Projection of the network activity on the first three PCs. The color code stands for response angle. At any time-within-trial, responses lie on a ‘ring’ encoding the response angle. **F** Projection onto different PCs. The manifold is not entirely contained in the space spanned by the first 3 PCs, but “stretched” along several orthogonal axes. **G** The multiscale ID plot shows ID estimates as a function of *K*, in black (reliable estimates) or red (unreliable ones). At high *K* > 500, all estimates become unreliable. **H** The local ID histogram shows a narrow peak around 2. Estimating the mode of the distribution yields *ID*_*FCI*_ = 2.2. A linear ID estimation method, parallel analysis (PA), finds *ID*_*PA*_ = 7. The classical geometric maximum-likelihood estimator (MLE) yields *ID*_*MLE*_ = 3.34, while Two-NN yields *ID*_*TwoNN*_ = 7.03, a severe overestimation due to its high sensitivity to local noise. **I** We used locally linear embedding (LLE) to find a global low-dimensional representation of the neural manifold for RT-Go. LLE flattens the manifold onto a circle spanned by a radial coordinate, representing time-within-trial, and an angular coordinate, representing response angle.

Before analyzing how the manifold’s ID varies across tasks and trainings, we provide a detailed analysis of an exemplar case: the neural manifold of a RNN performing the React-Go (RT-Go) task, where the network must respond with an output in the same direction of the input stimulus (Fig. 3B). We first performed principal component analysis (PCA). The first PC explains 64% of the variance, while four subsequent ones explain 5-10% (Fig.3C). As a result, parallel analysis (PA) finds *ID*_*PA*_ = 7. In the space defined by the first 3 PCs, the manifold appears to be topologically equivalent to a cone (Fig. 3D,E). The cone axis is roughly aligned to the first PC, while for fixed values of the first PC, the points describe a (deformed) circle in the space spanned by the second and third PC. In Fig. 3D, points are colored according to time-within-trial (*t* = 0 start of trial; *t* = 152 end of trial); in Fig. 3E, according to response angle. These representations allow an easy grasp of the manifold’s shape. Before the stimulus presentation, the network is silent (**r** = 0). This configuration corresponds to the vertex of the cone. As the stimulus is presented, the network reacts by reaching a set of attractor states that depend on the response angle and approximately describe a circle. These attractor points are not reached immediately, hence trajectories span a ‘conical’ surface. Importantly, the surface does not lie exactly in the linear space spanned by the first three PCs. In fact, the cone is ‘tilted’ in several dimensions. This is shown in Fig. 3F, where we replace the third PC with successive PCs. As a result, linear estimates of the ID end up to be considerably larger than 3.

In principle, the ‘conical’ surface could be described by an axial coordinate (encoding time-within-trial) and an angular one (encoding response angle). Thus, we would expect *ID* = 2. lFCI gave *ID*_*lFCI*_ = 2.05. A closer inspection through the multiscale ID plot (in 3G) shows that local estimates are peaked around 2 for sufficiently small neighborhoods (*K ≲* 1000), while they become much higher for larger neighborhoods, eventually reaching values in the range [10, 20]. Yet, estimates obtained for *K ≳* 1000 are not reliable, as diagnosed by a poor goodness-of-fit (*GoF* > 0.02) and/or curvature (*δ* > 3). As a result, the local ID histogram has a clear peak around 2 (Fig. 3H). The intrinsically 2-dimensional geometry of the manifold can be effectively captured using locally linear embedding (LLE, [65]), which yields an explicit 2-dimensional parametrization of the manifold. As shown in Fig. 3I, the manifold can be ‘flattened’ on a two dimensional surface parametrized by a radial (time) and an angular (response angle) coordinate. However, the classical maximum likelihood ID estimator, however, yields *ID*_*MLE*_ = 3.34, which considerably lower than PA, but larger than 2. The Two-NN estimator gives *ID*_*TwoNN*_ = 7.03. As is well known, the MLE estimate can be affected by the non-uniformity of the distribution, in this case probably causing an overestimate. The Two-NN estimate, in turn, is strongly biased by small-scale noise [45]. Only by reducing the density of the dataset through decimation (which reduces the effects of small-scale noise increasing the average distance between points), we obtained an estimate *ID*_*TwoNN*_ = 2.05 much more consistent with the lFCI one (Fig. S3).

In summary, even in this simple task, the neural manifold is highly curved and tilted in several dimensions. As a result, linear ID estimation methods considerably overestimate the dimension. On the contrary, lFCI yields an estimate that is congruent with the task structure and the possibility of parametrizing the manifold with 2 coordinates.

In Fig. 4, we show ID estimates obtained when training 10 RNNs to solve each of the tasks. For all tasks in the Go Family, we consistently obtain 2 ≤ *ID*_*lFCI*_ ≤ 2.5 (Fig.4D). Classical *MLE* gave ID estimates between 3 and 4 (Fig.4B). Two-NN gave larger and quite variable ID estimates, *ID*_*TwoNN*_ ∈ [5, 12] (Fig.4C). When applying decimation, which removes small-scale noise, we obtained estimates consistent with those of lFCI, i.e., around 2 (Fig. S3D,F). This result is consistent with the above analysis of the RT-Go task, showing that the manifold is essentially spanned by a circular and a temporal variable. Linear ID estimates by parallel analysis (PA) are in general much larger (up to *ID*_*PA*_ = 10) and strongly depend on the task and training instance (*ID*_*PA*_ ∈ [6, 11] for Go and Anti; *ID*_*PA*_ ∈ [3, 5] for Dly-Go and Dly-Anti; *ID*_*PA*_ = 7 for RT-Go and Rt-Anti). PA counts all directions on which the manifold has a non-vanishing projection, which broadly corresponds to the number of PCs explaining 90% of the variance (Fig. S3A). Another linear ID estimation method, the participation ratio (PR), effectively counts only the most relevant projections. PR yields indeed lower estimates than PA (in the range [1, 5], but estimates again strongly depend on the task and training instance (Fig. S3B). For the RT and Dly tasks, the PR often yields estimates close to 1, a direct consequence of the presence of a ‘dominant’ PC explaining a large fraction of the total variance (the cone ‘axis’ representing time-within-trial).

**FIG. 4:**
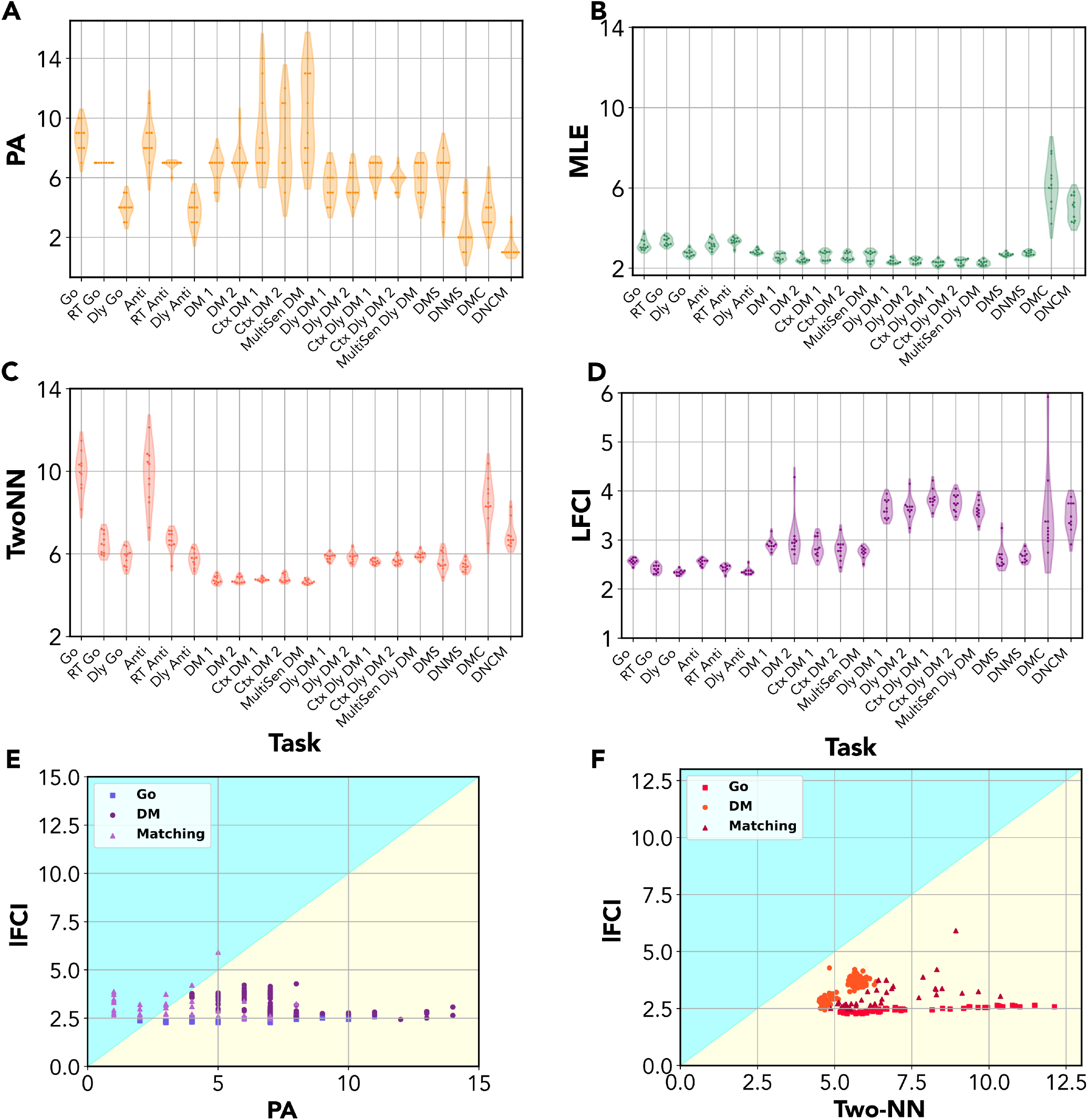
Intrinsic dimension of neural manifolds for the Cog-Task battery. For each task in the Cog-Tasks battery, we trained 10 independent recurrent neural networks (RNNs) and computed their intrinsic dimension (ID) with different methods. Notably, we compare estimates obtained by the local FCI (lFCI) method with those obtained with a linear ID estimation method, parallel analysis (PA) and two commonly used geometric ID methods, the classical maximum likelihood estimator (MLE) and the two-nearest-neighbors (Two-NN) estimator. **(A)** ID as the number of significant PCs identified with the parallel analysis (PA). For most tasks, ID estimates are not consistent in different trainings. Most estimates give *ID*_*PA*_ ≥5. **(B)** ID estimated by the classical maximum likelihood estimator (MLE). MLE gives very consistent estimates, with 2 ≤ *ID*_*MLE*_ ≤ 3 for most tasks. The only exception is given by the delayed-match-to-category (DMC) task, which, however, is very difficult to train, with most networks learning the task only approximately. **(C)** For tasks in the ‘Go-Anti’ family, Two-NN gives quite inconsistent estimates, with *ID*_*TwoNN*_ ≥ 6. For tasks in the ‘Decision Making (DM)’ family, estimates are consistent, with *ID*_*TwoNN*_ ≈ 5 for the non-delayed and *ID*_*TwoNN*_ ≈ 6 for the delayed tasks. For tasks in the ‘MS’ family, we obtain *ID*_*TwoNN*_ ≈ 5 for D(N)MS, and *ID*_*TwoNN*_ ≥ 6 for D(N)MC. **(D)** The local FCI (lFCI) method gives consistent estimates around 2.5 for all tasks in the ‘Go’ family, and estimates between 3 and 4 for tasks in the DM and MS family. **(E)** The ID identified by lFCI is always considerably lower than the ID estimate obtained by PA, except for the MS family. This is a reflection of the fact that neural manifolds for the Cog-Tasks battery are generally curved and extended across many linear dimensions, despite having an intrinsic 2- or 3-dimensional geometry. **(F)** The ID identified by FCI is always lower than the estimate obtained by Two-NN, which is severely affected by small-scale noise.

For tasks in the DM family, we consistently obtained 2.5 ≤ *ID*_*lFCI*_ ≤ 3 for the non-delayed tasks, and 3.5 ≤ *ID*_*lFCI*_ ≤ 4 for the delayed tasks. We can trace back this increased dimension to a perturbation of the previously analyzed ‘conical’ manifold structure. Like in the Go family, neural trajectories must encode the response angle in the ‘Go’ phase to produce a correct response. However, in the delayed tasks (where the two stimuli are given in succession) the network must also transiently encode the first stimulus angle, which must be kept in memory until the second stimulus is presented and the correct response can be computed. Trajectories converging to the same attractor points thus form a smeared bundle. This bundle also includes ‘backward’ trajectories: as the stimuli are switched off in the ‘Go’ phase, trajectories do not remain in the vicinity of attractor points but slowly converge back to the origin before the end of the trial (for a detailed example and discussion, see Appendix B and Fig. S2). This ‘smearing’ of trajectories associated to the same attractor determines an additional ‘transversal’ degree of freedom, causing a rise in dimension. This phenomenon also occurs in the non-delayed tasks, where ‘smearing’ is only due to backwards trajectories not coinciding with the forward ones, causing only a minor dimension increase with respect to the Go tasks (~ +0.5). ID Estimates obtained with classical *MLE* are consistent and slightly above 2 for all DM tasks (4B), suggesting that MLE identifies only the two local dimensions with the strongest variability. ID Estimates obtained with Two-NN are consistently larger than those of lFCI (*ID*_*Two*−*NN*_ ≃ 5 for the non-delayed DM tasks, *ID*_*Two*−*NN*_ ≃ 6 for the delayed ones; Fig. S3C). Upon decimation, estimates are in good agreement with those of lFCI (S3D). On the other hand, linear ID estimates are again quite inconsistent over tasks and trainings: PA yields generally much larger estimates (*ID*_*PA*_ ∈ [4, 15]; Fig. S3A), while PR yields estimates in a similar range (2 ≤ *ED*_*PR*_ ≤ 5). PR estimates are not correlated those of lFCI (Fig. S3E), suggesting that the two methods capture different geometrical properties: *ID*_*PR*_ captures global variation along different orthogonal axes, *ID*_*lFCI*_ captures local variation.

Finally, for tasks of the MS family, we consistently obtain *ID*_*lFCI*_ ~ 2.5 for DMS and DNMS. This result is fully consistent with estimates given by MLE and Two-NN (after decimation). PA gives inconsistent and high estimates, while PR gives estimates close to 1, due to a dominant PC. For the DMC and DNCM tasks, we obtain 2 ≤ *ID*_*lFCI*_ ~ 4 (except for a single training with *ID*_*lFCI*_ ~ 5). ID Estimates are much more variable than for all other tasks. This variability, which also emerges in MLE and Two-NN, probably reflects a difficulty of properly training these tasks (the accuracy was 0.90 ± 0.05 compared to 0.97 ± 0.01).

### C. ID of synthetic neural recordings

Altan et al. [31] introduced a benchmark data set of synthetic neural recordings. As detailed in Methods, the data were generated by considering firing rate data from *d* independent recording channels, with varying *d* ∈ [3, 6, 10, 20, 40], embedding them in *N* = 96 dimensions with a linear transformation (*linearly embedded data*), and finally applying a non-linear transformation (*nonlinearly embedded data*).

The linearly embedded data present a challenge for classical geometric ID estimators, but not linear ones. While PA always finds *ID* ≃ *d*, geometric estimators find *ID* ~10 for all *d* ≥ 10, a reflection of their (known) difficulty in estimating high dimensionalities. Here, lFCI outperforms geometric methods, giving estimates broadly in line with PA ones: *ID*_*lFCI*_ = [3.00, 5.63, 8.87, 16.74, 29.41] for *d* = [3, 6, 10, 20, 40] (Fig. 5B). In fact, this data do not pose significant challenges to lFCI. This is reflected in the very well-behaved appearance of the multiscale ID plots and the local ID histograms for *d* = 6, 20, 40 (Fig. 5C,E,F). Estimates are reliable (a natural consequence of the non-curved nature of the data) and consistent for all scales *K*, so that the local ID histogram is sharply peaked. The ID underestimation (which becomes evident for *d* = 20 and even more so for *d* = 40) is due to the presence of many dimensions with extremely small variance, *<* 10^−2^ ((Fig. 5A). In this condition, it is essentially impossible to reliably discriminate these dimensions from small-scale noise. In fact, also PA tends to identify a number of dimensions lower than *d*, and *ID*_*lFCI*_ and *ID*_*PA*_ converge for *d* = 40. To validate this hypothesis, we variance-normalized the data before the embedding. For these equal-variance data, we obtained *ID*_*lFCI*_ ≃ *d* for all *d*.

**FIG. 5:**
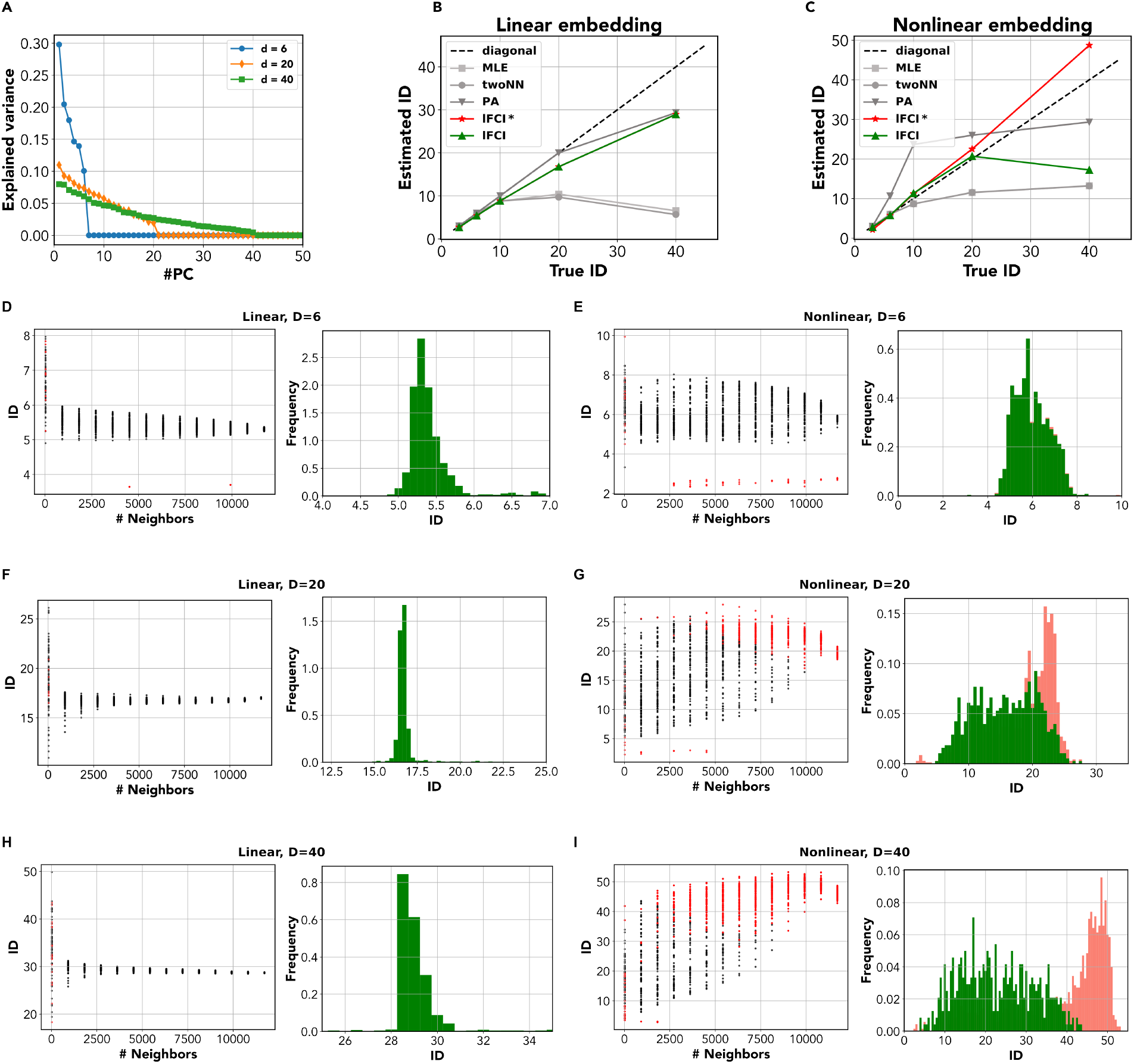
ID estimation for the synthetic neural data data with increasing dimension. In Ref. [31], neural manifolds were constructed by randomly sampling firing rates from *d* independent channels in a multielectrode array. Data were then linearly embedded in 96 dimensions (*linearly embedded data*), and further subjected to a non-linear transformation (*nonlinearly embedded data*). **A** Variance explained by the principal components (PCs) of the linearly embedded data. **B** The ID was estimated by different methods, including parallel analysis (PA), Maximum Likelihood Estimator (MLE), Two-Nearest Neighbors (Two-NN), the standard global FCI method, and the local FCI method. Dashed line indicates the identity. The linearly embedded data are stretched along *d* orthogonal axes, but the variance is very small (≲ 10^−2^) for several components. All ID estimation methods yield *ID* ~*d* for *d* ≤ 10. For *d* ≥10, MLE and Two-NN show a non-monotonic behavior, with *ID* ≤ 10. PA, FCI and Lci give monotonically increasing estimates for increasing *d*, with *ID*_*lFCI*_ ~*ID*_*TwoNN*_ ~ *ID*_*FCI*_ ~30. **C** The nonlinearly embedded data give rise to strongly non-smooth manifolds. All ID estimation methods yield *ID* ~*d* for *d*≤ 10. For *d* ≥ 10, MLE and Two-NN show a non-monotonic behavior, with *ID* ≤ 10. lFCI show a monotonically increasing trend, but the ID remains much lower than *d* (*ID*_*FCI*_ ≤ 13). The global FCI gives *ID*_*lFCI*_ ~ *d* for all *d*. **D** For the linearly embedded data at *d* = 6, local ID estimates are consistent at all *K*, with a slightly decreasing trend for increasing *K*, and we obtain *lFCI* = 5.63. **E** For the nonlinearly embedded data at *d* = 6, local ID estimates are consistent at all *K*, and we obtain *lFCI* = 5.8. **F** For the linearly embedded data at *d* = 20, most local ID estimates are consistently in the range [15, 20] at all *K*, with a decreasing trend for increasing *K*. We obtain *lFCI* = 16.74, consistent with the number of PC components with variance > 10^−2^. **G** For the nonlinearly embedded data at *d* = 20, local ID estimates are very variable (in the range [5, 20] for all *K <* 10^4^. Only for *K* ≈ 10^4^ do estimates become more consistent with values around 20. **H** For the linearly embedded data at *d* = 40, most local ID estimates are consistently in the range [25, 35] at all *K*, with a decreasing trend for increasing *K*. We obtain *lFCI* = 29.2, consistent with the number of PC components with variance > 10^−2^. **I** For the nonlinearly embedded data at *d* = 40, local ID estimates are very variable (in the range [5, 40] for all *K <* 10^4^. Only for *K* ≈ 10^4^ do estimates become more consistent with values around 40.

The nonlinearly embedded data have some very peculiar characteristics, which we resume here (only some were discussed in [31]): i) the data distribution along each dimension is nearly log-normal. ii) The upper tails of the distributions give rise to isolated low-density regions along orthogonal axes (Fig. S4A). In these regions, the ‘tangent plane’ has not a dimension equal to the ID, and perhaps is not even well-defined. iii) the density varies dramatically across the dataset (to measure this quantitatively, we used the inverse of the distance of the 10-th neighbor as a proxy of the density). As a result, the manifold contains a highly dense ‘core’ and a large, rarefied ‘periphery’. To better visualize this, in Fig. S4B,D we projected the manifolds for *d* = 20, *d* = 40 in two dimensions using t-stochastic neighbor embedding (TSNE) [66], showing the local point density.

Due to these joint characteristics, ID estimation becomes a difficult endeavour. Due to i), the covariance structure of the data is strongly influenced by outliers: this makes linear ID estimation unreliable. In fact, PA first overestimates (for *d <* 20), then underestimates (for *d* = 40) the ID. Geometric estimators are bound to underestimate the ID (for *d* > 10) due to their intrinsic limitations but also the manifold geometry, as nearest-neighbor statistics in the peripheral regions strongly deviates from that predicted shape. Indeed, MLE and Two-NN give *ID*_*MLE*_, *ID*_*Two*−*NN*_ *< d* for *d* 10. In this situation, lFCI can give accurate estimates up to *d* = 20, while it fails for *d* = 40, producing a sharp underestimation: *ID*_*lFCI*_ = [2.8, 5.8, 11.3, 20.7, 17.2] for *d* = [3, 6, 10, 20, 40]. The problem for *d* = 40 can be mitigated by relaxing a step in the pipeline, the removal of estimates corresponding to bad GoF, although this comes at the price of some overestimation: *ID*_*lFCI*_ = [2.5, 5.8, 11.2, 22.5, 48.7] for *d* = [3, 6, 10, 20, 40]. To better analyze the reasons for this (partial) success, in Fig. 5E,G,I we show the multiscale ID plots and the local ID histograms for *d* = 6, 20, 40. For small *d* (*d* = 6), the behavior is similar to the one observed in the linear embedding case. Although local ID estimates are slightly more variable (in the range [5, 7]), the local ID histogram is still sharply peaked For *d* = 20, 40 local ID estimates appear to be highly variable for most neighborhood sizes *K*, except *K* ≥ 10000. This behavior depends on the dataset geometry: local ID estimates strongly correlate with the density (Fig. S4C,E). Neighborhoods in the highly dense ‘core’ yield local ID estimates ~*d*, whereas neighborhoods in the rarefied periphery cannot detect the full dimensionality, systematically yielding lower ID estimates. For large *K* (≳ 5000), all neighborhoods encompass a large fraction of the whole manifold. ID estimates become accordingly more consistent. However, estimates also become unreliable. This is not due to curvature (we always find *δ* ~ 1 in our data, reflecting the absence of curvature), but rather to a bad GoF. In fact, large neighborhood must include regions with variable density, leading to a violation of the isotropy assumption and hence a deviation of the FCI curve from its theoretical shape. The FCI fitting procedure, however, spots a reasonable if slightly overestimated dimension. For *d* = 20, the local ID histogram has a peak around 20 even if unreliable estimates are discarded, even though the peak is shifted to ~22 and becomes more pronounced if they are included (5G). For *d* = 40, the local ID histogram has a peak around 20 unless unreliable estimates are included (5G), which produces a peak around 45. In synthesis: when considering small neighborhoods, the presence of rarefied regions tends to produce ID underestimation; when using large neighborhoods, this problem is solved but density variations imply a bad GoF and some overestimation. Nevertheless, even for this very challenging manifold, lFCI can yield rich insight into the manifold’s structure and ID.

## IV. DISCUSSION

The dimensionality of neural population activity is at the root of key open issues in computational neuroscience, such as the relation between the neural activity and the structural architecture of neural circuits [17, 19, 67], the relation between spontaneous and task-evoked activity [68–70], and the development of abstract codes [23, 26, 71]. More in general, characterizing the dimensionality of neural manifolds is critical to understand how neuronal populations collectively encode relevant variables and perform computation [7]. However, this is currently a matter of wide debate, given the abundance of conflicting findings of low-vs. high-dimensional neural activity. Classic examples are the seminal study by Churchland et al. demonstrating that a reaching task induces a low-dimensional neural activity [14] and the article by Stringer et al. showing an unbounded scaling of dimensionality with increasing number of recorded neurons in visual cortex [68]).

### Methodology

The ongoing discussion would be highly benefited by the development of robust methodologies to measure the intrinsic dimension (ID) of neural manifolds [16]. Linear methods (PCA and its evolutions) fail in presence of curvature and nonlinearity, and are thus inappropriate in the (common) case where neural manifolds are strongly nonlinear [41]. Geometric ID estimators (which have been extensively applied to study representations of deep artificial neural networks [72–75]) might provide a valid alternative, but they tend to be fragile to undersampling [31]. This is a major impairment, leading to dramatic underestimation of the ID in the case of short recordings and/or high dimensionality. Thus, currently widespread ID estimation methods are not consistently reliable within the broad landscape of neural manifolds.

Here, we introduced a novel ID estimation pipeline, based on a local version of the full-correlation integral (lFCI). Our pipeline computes the ID locally on neighborhoods of variable size, identifying neighborhoods that deviate from the local tangent plane or have a non-isotropic distribution, for which local ID estimates are unreliable. Eventually, a robust global ID estimate can be obtained from the histogram of local ID estimates. Our pipeline is versatile enough to reliably cover a wide range of dimensionalities and non-linearities, as shown by the result summary in Table I. In all cases analyzed, lFCI performs comparably or better than other methods. Importantly, no other method (among those included in our analysis) was able to yield consistently reliable results in all cases. Our pipeline encountered tangible difficulties only in the extreme case of a high-dimensional, highly skewed data distribution; even in this case, relaxing the pipeline yielded correct insight on the ID.

**TABLE I:** Summary of the results.

| Task name | True ID | PA | MLE | twoNN | IFCI |
| --- | --- | --- | --- | --- | --- |
| cog-task (GO) | $\sim 2$ | 6.43 | 3.09 | 7.42 | 2.44 |
| cog-task (DM) | $\sim 3$ | 7.08 | 2.23 | 5.26 | 3.29 |
| cog-task (Match) | $\sim 3$ | 3.40 | 4.14 | 6.63 | 3.08 |
| neural traj. (lin) | 3 | 3.00 | 3.04 | 2.99 | 2.70 |
|  | 6 | 6.00 | 5.94 | 5.88 | 5.40 |
|  | 10 | 10.00 | 8.82 | 8.77 | 8.84 |
|  | 20 | 20.00 | 9.70 | 10.40 | 16.77 |
|  | 40 | 29.33 | 5.66 | 6.54 | 28.94 |
| neural traj. (exp) | 3 | 3.00 | 2.98 | 3.02 | 2.80 (2.13*) |
|  | 6 | 10.67 | 6.10 | 6.04 | 5.77 (5.77*) |
|  | 10 | 23.67 | 8.65 | 8.69 | 11.37 (11.20*) |
|  | 20 | 26.00 | 11.58 | 11.52 | 20.71 (22.51*) |
|  | 40 | 29.33 | 13.20 | 13.24 | 17.24 (48.72*) |

In future developments, we might integrate this pipeline with suitable denoising procedures to further enhance its robustness. In Ref. [31], Altan et al. proposed the following denoising/estimation scheme: 1) use parallel analysis (PA) to obtain an upper bound on the ID, *D*_*u*_ 2) denoise the data by first projecting (linearly, via PCA, on non-linearly, via an autoencoder) onto a *D*_*u*_-dimensional space, and then projecting back into the original space 3) if nonlinear denoising is more effective than linear denoising (indicating a nonlinear manifold) use a geometric ID estimator; otherwise, use a linear one. Our results show that FCI can yield a more robust upper bound than PA, also valid in a highly nonlinear case. Moreover, we showed that linear methods do not provide any specific advantage over FCI. We thus suggest the following modification of Altan et al.’s scheme: 1) use FCI to obtain an upper bound on the ID, *D*_*u*_ 2) denoise the data 3) characterize the ID using lFCI.

### Low-dimensional neural manifolds for Cog-Tasks

Recent literature has abundantly mentioned ‘low-dimensional neural manifolds’. Within experimental studies, the low-dimensionality was generally inferred from the fact that few PCs could explain a large fraction of variance. However, only rarely was a rigorous characterization of ID performed (an exception is [16]). Theoretical studies have often investigated cases where neural activity was *compelled* to be low-dimensional by virtue of structural constraints, such as a low-rank connectivity [17, 18, 59]. To what extent networks *trained* to perform simple task would exhibit a low dimensional activity was therefore an open question. In this work, we have analyzed the activity of RNNs performing the common ‘Cog-Task’ battery, showing that their activity is indeed organized in low-dimensional (approximately 2- or 3-dimensional) manifolds. The low-dimensionality seems to be directly related to the simplicity of the tasks at hand. The only key variable the networks must encode is the response angle - a relatively simple function of the angular inputs which can be computed even by a network of non-connected units with memory [49]. Response angle can be encoded by a ‘ring’ of attractors, which can be reached in a finite time by trajectories starting from the origin. This structure implies a minimal dimensionality of 2 for the neural manifolds. In some tasks, the response angle only determines the endpoint of the trajectories, but not the whole path, an additional variability reflected in a dimension increase of ~ +1. In future studies, we could investigate more complex tasks such as the ‘Mod-Cog’ battery proposed by [49].

### ‘Embedding’ vs. ‘intrinsic’ dimension of neural manifolds

In Ref. [40], Jayazeri et al. explicitly distinguished between the ID and the minimum dimension of a linear subspace containing the manifold, which they termed ‘embedding dimension’ (ED). Following their terminology, linear methods (such as parallel analysis, participation ratio, and PCA with a fixed variance threshold) essentially measure the ED. Many research works simply employ linear methods without paying attention to the ID/ED distinction (see, e.g., [13, 14, 19, 22, 26, 27, 32–36, 38]), and many recently proposed developments for dimensionality reduction are also based on linear projections [24, 33, 37, 39]. However, since neural manifolds can be highly non-linear and curved [41], the ID and the ED may considerably differ. A paradigmatic example is provided by our analysis of neural manifolds of RNNs trained to perform the Cog-task battery. Albeit intrinsically low-dimensional, these manifolds had non-vanishing projections along several orthogonal axes, and the ED was considerably higher than the ID. Importantly, the ID, but not the ED, was shown to be fundamentally related to encoding and computation. In the ‘Go family’, all trained networks displayed an ID close to 2, while the ED was contingent on the specific instance of trained network analyzed: for the same task, it could vary in a wide range (from 6 to 11). In the DM task, the ID was close to 3 for all trained manifolds, while PA led to a wide range of EDs, in the range [3, 15], with large fluctuations for the same task (e.g., from 6 to 15). Our findings show that fundamental task variables such as response angle and integration time can be represented *non-linearly* within the manifold, even in simple cases, and may not be easily captured in terms of principal components. The weak dependence of the ID on the training instance suggests that the ID may be among the ‘universal’ properties [76] that do not depend on network architecture and training, but only on the computational scaffold underlying the task [76]. In a classic paper, Gao et al. proposed an upper bound to the neural manifold dimension in terms of the ‘neuronal task complexity’ (NTC) [36], an index depending on the autocorrelation of activity for similar values of task-relevant variables (such as response angle and integration time). In fact, the NTC bound applies to a measure of ED, the participation ratio (PR), and this bound is generally quite loose and much larger than the ID (Fig. S3). Finally, a technical weakness affecting most ED estimators is their inherent dependence on a threshold to define the ‘relevance’ of principal components (PA is no exception, as it depends on an arbitrary significance threshold). The PR does not have this limitation, but it is strongly biased when variance along several PC is highly unequal. As observed for the Go and the MS family tasks, as the PC associated with time had a much larger variance than the others, we obtained an underestimation with *ID*_*PR*_ ~ 1. Such an estimate completely neglects the (crucial) angular variable.

### Limitations

The main limitation of the lFCI method is that it assumes a smooth manifold structure. In non-smooth, sparsely or radically non-uniformly sampled, local neighborhoods may deviate from the concept of a tangent plane. Correspondingly, some of the dimensions may be poorly sampled locally, yielding local ID estimates that are significantly lower than the global ID of the manifold. We observed this phenomenon in the analysis of the nonlinear data from [31], where low-density, peripheral regions yielded a lower local ID than the high-density bulk of the manifold. These nonlinear data were obtained by artificially applying a non-linear transformation to real data sampled from independent multi-electrode recordings. The nonlinearity of the ensuing data does not result from curvature, but rather from strongly skewed, fat-tailed firing rate distributions. As we do not know whether similar distributions are likely to arise in real cases, future analyses may focus on experimental data from large-scale recordings to test for the distribution of firing rates. LFCI was able to yield insight in the ID and manifold structure even in this extreme situation, but a careful analysis was required. We do not currently know to what extent FCI would still work in cases where its core assumptions are completely violated, such as non-uniform binary data or radically large (> 1000) dimensions where the very concept of ‘manifold’ may start to falter. In these situations, completely different methods may be required, departing from common linear and geometric ID estimators [77].

### Conclusion

In conclusion, we proposed a robust pipeline to estimate the intrinsic dimension of neural manifolds, offering a new tool to tackle an open debate. The inherent properties of lFCI make it robust to undersampling and nonlinearity, allowing us to probe a large range of IDs. Our pipeline allowed us to rigorously demonstrate that simple tasks induce a low-dimensional structure in neural activity and highlight the difference between the ID and the ‘embedding dimension’ given by linear estimators.

## V. ACKNOWLEDGMENTS

This work recevied financial support form the Italian Ministry of University, PRIN grant 2022HSKLK9, “Unveiling the role of low dimensional activity manifolds in biological and artificial neural networks”, CUP C53D23000740006/, CUP D53D23002220001. We thank Alessandro Laio for precious suggestions, and Luca Mazzucato, Samir Suweis and Bill Bialek for their feedback on the paper.

## VI. DATA AND CODE AVAILABILITY

The implementation of the lFCI method and notebooks reproducing some of the results in this paper are available on GitHub, https://github.com/JFadanni/ExploringNeuralManifolds.git.

---

**FIG. S1:**
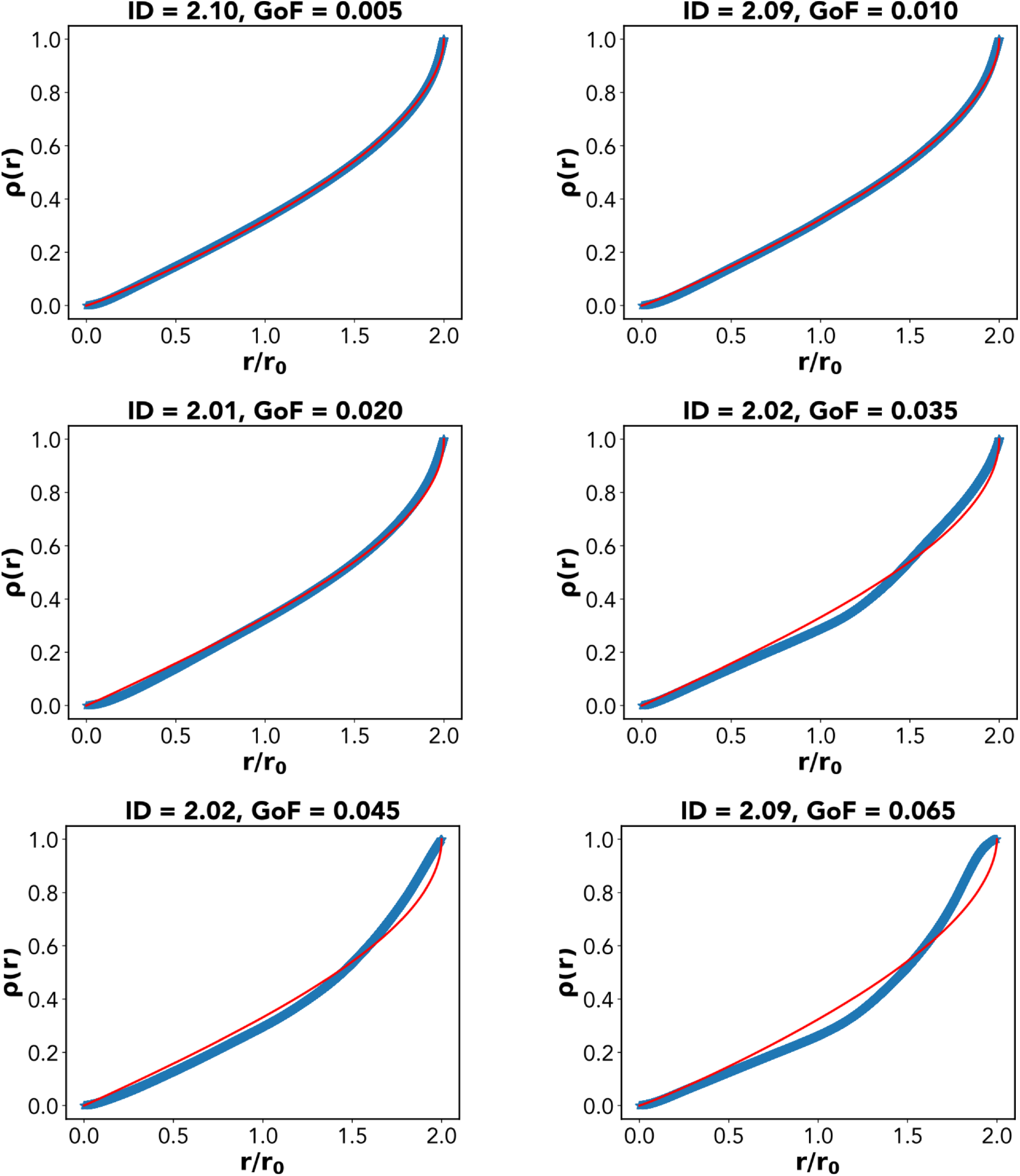
Examples of Goodness-of-Fit (GoF) for FCI

## VIII. SUPPLEMENTARY MATERIALS

### A. Goodness-of-Fit for FCI

### B. Neural manifold for Ctx-DM1

In the Ctx-DM1 task, the network must ignore the stimuli presented in modality 2 (**u**_*mod*2_, and respond in the direction of the stronger of two stimuli presented in modality 1 (**u**_*mod*1_, as soon as the fixation cue goes off (S2A). Thus, the task is conceptually very similar to Fd-Go, except that the response angle is a more complex function of the input (the network should also ignore any input in modality 2, but this requirement can be trivially satisfied by having weak input connections from inputs units corresponding to modality 2 and recurrent units). Another difference is that the stimuli are turned off as soon as the ‘Go’ signal starts, which, as we shall see, has a relevant effect on the structure of the neural manifold.

In Fig. S2B we display snapshots of neural trajectories at different values of time-within-trial, coloring the points according to response angle. Before stimulus presentation, the RNN activity is at the origin (**r** = 0. Stimulus presentation determines a weak displacement from the origin. After the fixation cue goes off, activity moves towards a set of attractors representing different response angles. Similar to the case of RT-Go, these attractors are arranged onto an approximately ring-like structure on which response angle is represented as a circular variable. However, the structure of this ring is much less regular than in the RT-Go case. Because input ceases as soon as the fixation cue goes off, after reaching this ‘ring’, trajectories slowly go back towards the origin. Importantly, the ‘backward’ trajectories are not fully equivalent to the ‘forward’ trajectories (S2D), creating transversal additional variation along a direction orthogonal to the ones encoding for angle and time-within-trial (conferring a sort of ‘thickness’ to the surface). When visualizing the entire manifold (S2E), a highly twisted conical surface emerges. In Fig. S2C, we plot the percentage of variance explained by several principal components. The first two components explain, respectively, 24% and 20% of the variance; the five subsequent components explain between 5% and 12% each. As a consequence, a linear ID estimation method, parallel analysis (PA) gives *ID*_*PA*_ = 7. We estimated the ID with the lFCI. The multiscale ID plot (S2)F) shows that local ID estimates for *K* ≥ 10^3^ are increasingly unreliable (worse GoF). Estimates for *K* ≤ 10^3^ are mostly in the range [2, 5]. The local ID histogram (S2G) shows a broad peak between 3 and 4, giving *ID*_*lFCI*_ = 3.5. Consistently, MLE yields *ID*_*MLE*_ = 3.34. The Two-NN estimator gives a much larger estimate *ID*_*Two*−*NN*_ = 7.03, probably a consequence of small-scale noise. When using a multiscale version of Two-NN, we obtain indeed *ID*_*Two*−*NN*_ = 3.78.

We tried to project the manifold in two dimensions using locally linear embedding (S2H). LLE can only approximately ‘flatten’ the manifold onto a two-dimensional ring where the radial coordinate represents time-within-trial, and the angular coordinate represents response angle. Yet, some of the response angles are badly represented, probably because of the inability of LLE to separate the backward and forward trajectories.

### C. Intrinsic dimension of neural manifolds for the Cog-Task battery - Additional Results

In Fig. S3A, we show ID estimates obtained by counting the number of principal components (PCs) explaining 90% of the variance. These estimates are generally in line with estimates given by PA.

In Fig. S3B, we show ID estimates obtained by the participation ratio (PR), which effectively counts only the most relevant PCs. For tasks in the Go family, PR yields yields lower estimates than PA (in the range [1, 5], with estimates depending on the training instance. We obtain *ID*_*PR*_ = 3.5 ± 0.9 and *ID*_*PR*_ = 3.4± 0.8 for Go and Anti; *ID*_*PR*_ = 1.9 ± 0.4 and *ID*_*PR*_ = 1.9 ± 0.4 for Dly-Go and Dly-Anti; *ID*_*PR*_ = 1.13 ± 0.08 and *ID*_*PR*_ = 1.1 ±0.1 for RT-Go and Rt-Anti). For the RT and Dly tasks, the PR often yields estimates close to 1 (1≤*ED*_*PR*_ ≤ 2). This is a direct consequence of the presence of a ‘dominant’ PC explaining a large fraction of the total variance. While the manifold is certainly stretched across this direction (the cone ‘axis’ representing time-within-trial), it cannot be really embedded in a one-dimensional space, lest the angular information (which is essential to solve the task) be completely lost. For tasks in the DM family, PR yields 2 ≤ *ID*_*PR*_ ≤ 5, with variable estimates depending on the training instance. This suggests that, when considering only the strongest PCs, the manifolds are contained within a linear subspace of dimension ≤ 5. For tasks in the MS family, PR yields estimates close to 1, again reflecting the presence of a single dominant PC. Notably, while the range of values is somewhat similar, *ID*_*PR*_ and *ID*_*lFCI*_ are not correlated (Fig. S3E), implying that they capture different geometrical properties: the dimensionality of *ID*_*PR*_ captures global variation along different orthogonal axes, The *ID*_*lFCI*_ captures local variation.

In Fig. S3C we show the results of computing the ‘neuronal task complexity’ (NTC) metric. Gao et al. [35, 36] proved with a general theoretical argument that NTC is an *upper bound* on the linear dimensionality of neural manifolds given by PR.

**FIG. S2:**
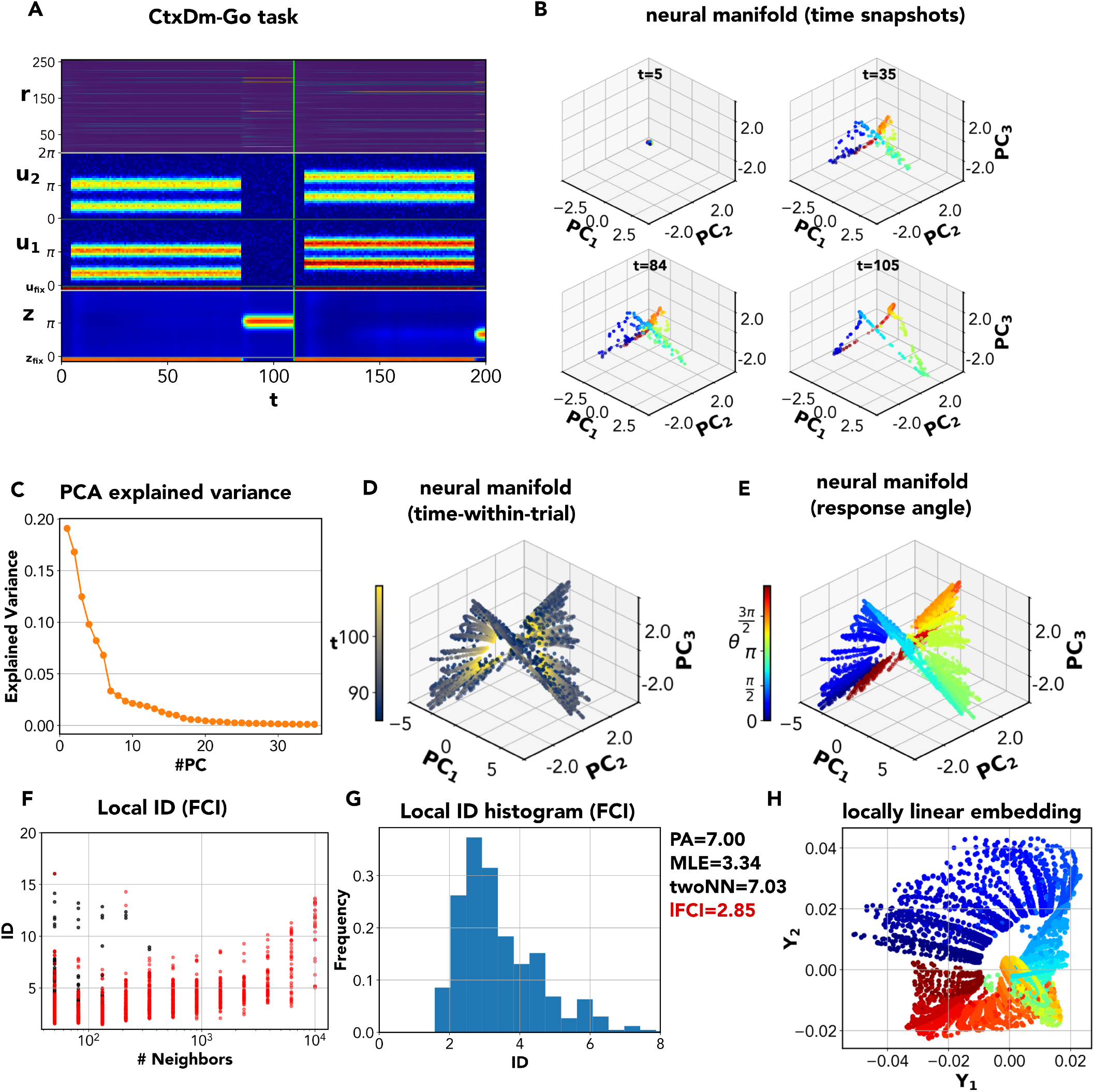
Dimensionality of the neural activity manifold in the Ctx-DM1 task. **A** Input, output and network activity for the Ctx-DM1 task; **B** Input, output and network activity for the Ctx-DM1 task; **C** Explained variance of the different PCs of the network activity. Looking at the variance, PCA predicts a dimensionality of 1 if we consider the highest drop in variance or a dimension of 7 if we consider the second elbow in variance. **D** Projection of the network activity on the first three PCs. The color code stands for time-within-trial (time form trial onset). *PC*_1_ is nearly aligned with time-within-trial. **E** Projection of the network activity on the first three PCs. The color code stands for response angle. axes. **F** Multiscale ID plot for the RT-Go task. At low *K*, estimates are close to 3. **G** Local ID histogram, showing a peak at *D* = 2.85. **H** Locally linear embedding can flattens the manifold for the CTx-DM1 task onto a 2-D surface but only with some approximation.

In Fig. S3D, we show the results of using the multiscale version of the two-nearest-neighbor method. According to [45], multiscale ID estimates can be obtained by progressively ‘decimating’ the dataset, randomly selecting 100 *·* 2^*K*^ points, with *K* = 1, …, log_2_(*P/*100) and estimating the ID only on the selected points. For each *K*, the random choice can be repeated *M* times, and the average of the *M* estimates is kept as an estimate of the ID at the scale *K*. Note that the logic is similar to that of the multiscale FCI method, but the role of *K* is reversed. Two-NN is intrinsically local, as it considers the distances of the first two neighbors from each point. When using the whole dataset, we are considering the data at the *finest* spatial scale, while small values of *K* correspond to *coarse* spatial scales - the opposite of what happens for lFCI. As a robust ID estimate, one can consider the coarsest ID obtained with significant decimation of the points (*K* = 0), as estimates obtained for large *K* are often very sensitive to small-scale noise, which tends to inflate ID estimates [45].

### D. ID of multi-electrode neural recordings - Additional Results

**FIG. S3:**
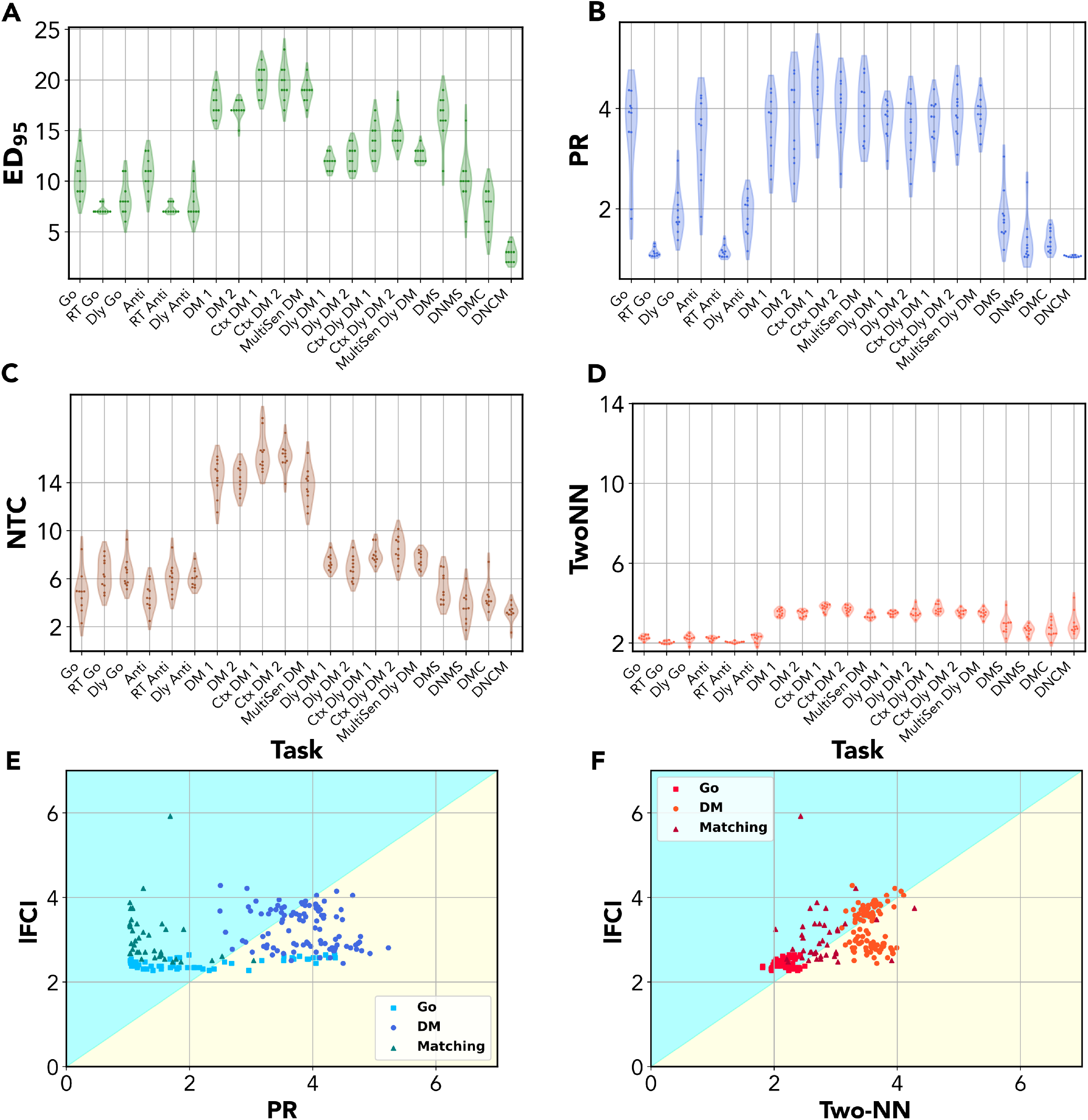
Intrinsic dimension of neural manifolds for the Cog-Task battery. For each task in the Cog-Tasks battery, we trained 10 independent RNNs to solve the task, and computed the intrinsic dimension (ID) of the neural activity manifolds for each case. **(A)** ID as the number of PCs identified with the principal component analysis (PCA) keeping the first principal components explaining 90% of the variance **(B)** ID estimated by the the participation ratio (PR) **(C)** upper bound on the ID given by neuronal task complexity (NTC) **(D)** ID computed though th(PCA) keeping the first principal components explaining 90**(E)** NTC can under- and overestimate the ID **(F)** The ID identified by the local Two-NN method and the local FCI are in good agreement.

**FIG. S4:**
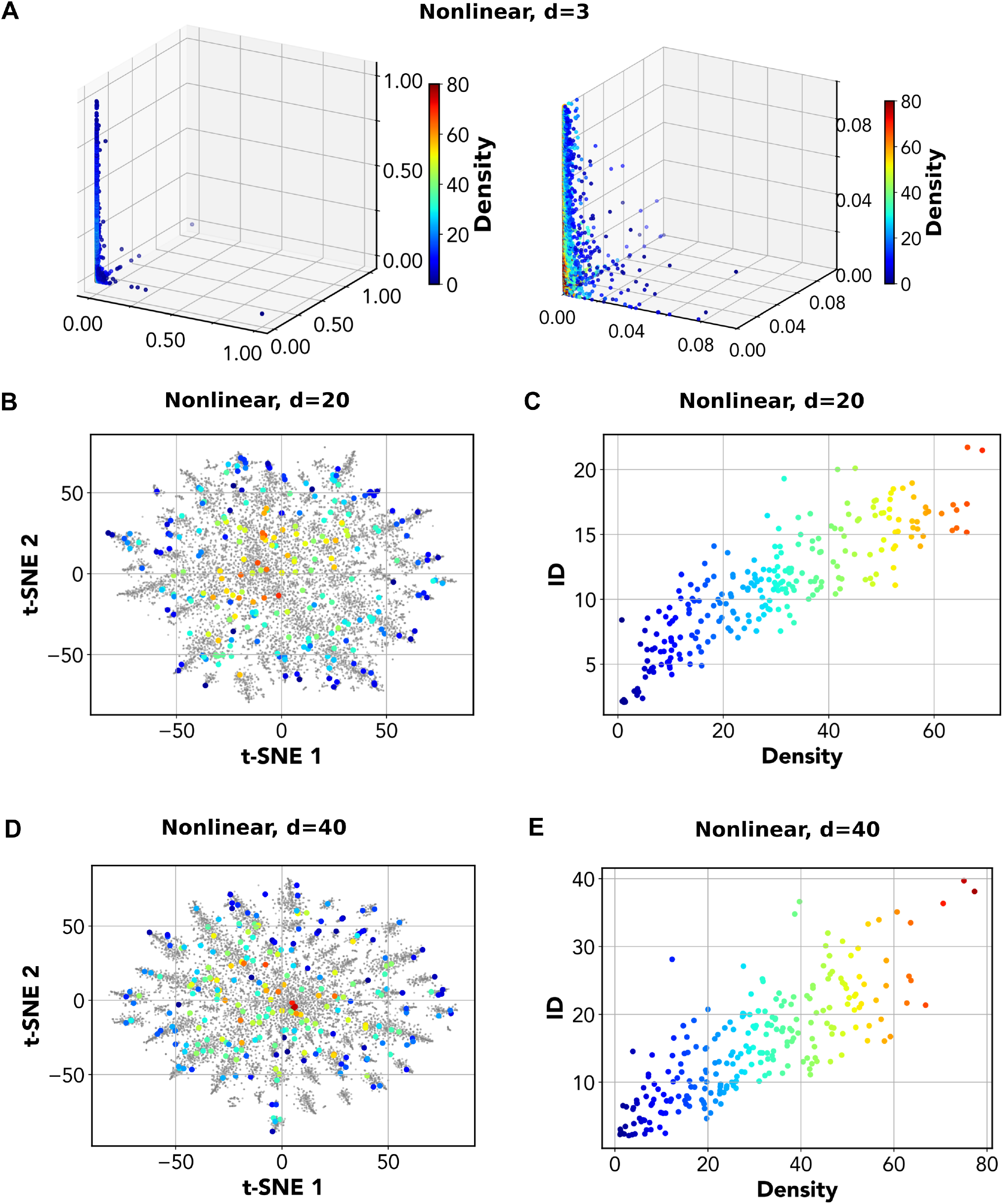
Local ID estimates and density for the ‘multielectrode array’ data. **(A)** TSNE embedding of the data with *d* = 20. Low local IDs are found in the ‘periphery’, while high local IDs are found in the ‘core’ **(B)** local IDs estimates strongly correlate with the local density of points for *d* = 20. **(C)** same as (A) for *d* = 40. **(D)** local IDs estimates strongly correlate with the local density of points for *d* = 20

**FIG. S5:**
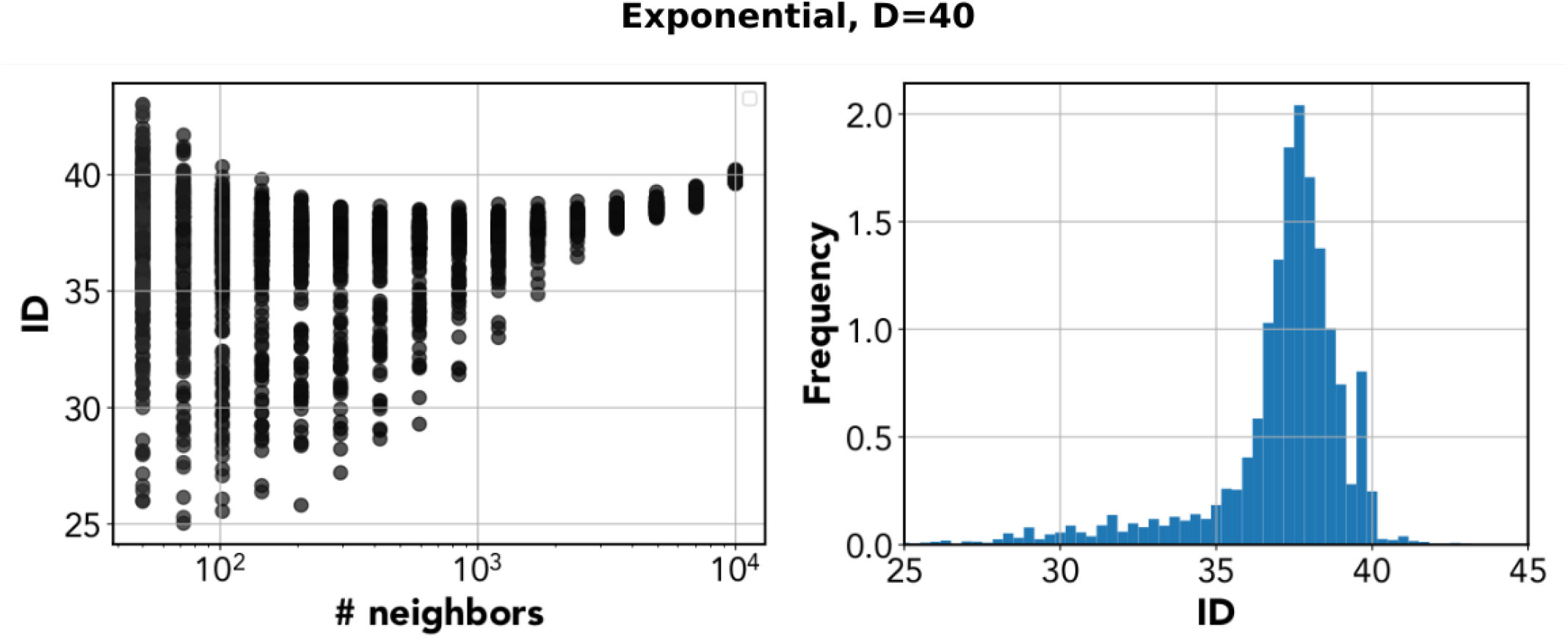
ID estimation for the ‘multielectrode array’ data with equal variance.

